# Winter rye cover cropping changes squash (*Cucurbita pepo*) phyllosphere microbiota and reduces *Pseudomonas syringae* symptoms

**DOI:** 10.1101/2021.03.10.434846

**Authors:** Rémi Maglione, Marie Ciotola, Mélanie Cadieux, Vicky Toussaint, Martin Laforest, Steven W. Kembel

## Abstract

Cover cropping is a soil conservation practice that may reduce the impacts of the economically important pathogen *Pseudomonas syringae* on crops including squash (*Cucurbita pepo*). To date, no studies have directly quantified the effect of rye cover crops on *P. syringae* populations, nor on the bacterial community of squash leaves. In this work, we tested the hypothesis that the protective effects of cover cropping on squash may be mediated by cover cropping effects on the plant’s microbiota that in turn protects against *P. syringae*. Using combined 16S sequencing and culture-based approaches, we showed that rye cover cropping protects squash against *P. syringae*, by decreasing pathogen population size on squash leaves and increasing fruit health and marketability at harvest. We also found evidence of a strong effect of rye cover crops on bacterial communities of the squash phyllosphere. Those findings were more striking early in the growing season. Finally, we identified numerous phyllosphere bacteria belonging to the genera *Sphingomonas, Methylobacterium* and *Pseudomonas* that were promoted by rye cover crops. Overall, our findings suggest cover cropping is effective for the sustainable management of *P. syringae* on squash and may provide a reservoir of potential microbial biocontrol agents colonizing the phyllosphere.

## Introduction

Cover cropping, or the growth of a plant to cover the soil for environmental benefits rather than for its harvest, is an increasingly popular option available to farmers to address the environmental and human health challenges associated with agricultural intensification (1). Cover cropping allows equivalent yield (2) or an increase in yield (3–5), weed control (6), nematode control (7), and reduces soil erosion (5,8). Winter cover crops used in northern countries are a promising avenue to reduce soil erosion and depletion by covering the soil during the winter (8). Cover crops can also improve water quality by reducing herbicide runoffs (9), and improve soil condition by reducing temperature variations and water loss (10). Cover cropping is known to shape the soil microbiome (11), but to date no study has quantified cover crop effects on aboveground microbial communities.

The phyllosphere microbiome, the microbial communities of aboveground plant parts, particularly leaves, are composed of a broad range of microorganisms such as bacteria, viruses, fungi and archaea (12). The microbiota on the aboveground parts of plants can improve plant fitness and biomass, primarily by reducing pathogen symptoms thanks to direct competition or associated with plant volatile compound (13,14). Microorganisms are also important pathogens of the phyllosphere: 20–30% of crop production losses worldwide are due to various pests and pathogens (15), and microscopic pathogens are estimated to account for 16% of losses (16). *Pseudomonas syringae*, one of the most widely studied bacterial plant pathogens, can infect a wide range of host plants including many economically important crops: it begins life as a leaf epiphyte and colonizes the host apoplast through wounds and stomates (reviewed in 17 and 18). Long-term intensive and frequent monocropping favour emergence of local pathogenic *P. syringae* reservoir (19). Efforts have been made to biologically control this pathogen on leaves using microbial competition: Linderman et al. (1987) used a competition population of *P. syringae* disarmed with an ice nucleation mutation to prevent pathogen-related frost damage on strawberry plants and Innerebner et al. (2011) used several *Sphingomonas* species on leaf surfaces to protect *Arabidopsis thaliana* plants from *P. syringae*. Moreover, many studies of plant induced systemic resistance (ISR) have noted potential leaf control of *P. syringae* based on the microbe-associated molecular patterns (MAMPs) mechanism and, interestingly, via microbial competitors located in the root-associated microbiome (reviewed in 22 and 23). For instance, Hossain et al. (2008) applied plant growth-promoting fungi *Penicillium sp*., isolated from soil, to promote tomato ISR against the leaf pathogen *P. syringae* pv. *tomato* DC3000.

Despite the potential for microbiota-based biological control of *P. syringae*, management of this pathogen is primarily done through copper applications, which has led to the development of resistance (25). Past studies have suggested that cover crops may provide soil borne biological control of pathogens (26–28), and we have found that rye cover crops helped to reduce *P. syringae* symptoms incidence on squash leaves (Toussaint et al., personal communication). The mechanism of this protective effect of rye cover crops against squash bacterial leaf spot is not known, but we hypothesize that it may be mediated by cover cropping effects on the plant’s microbiota that in turn protects against *P. syringae*.

In this study we used sequence and culture-based approaches to quantify the effects of different cover cropping approaches on bacterial communities on squash leaves infected by *P. syringae*. We first evaluated if cover crops could help to reduce *P. syringae* populations on squash leaf surfaces by direct measurement of pathogen abundance on leaves. We also considered the fruit’s health and marketability at harvest in such cropping practices. We then estimated the effects of different cover cropping methods on phyllosphere bacterial communities by quantifying leaf microbiome diversity and composition using a bacterial metabarcoding approach. Finally, we identified the bacterial taxa that were most strongly influenced by cover cropping practices.

## Material &Methods

### Experimental design and field treatment

All samples in this study were collected from the Agriculture and Agri-Food Canada L’Acadie Experimental Farm at Saint-Jean-sur-Richelieu, Quebec, Canada (45°17’48.7”N 73°20’14.8”W). The experiment comprised 6 replicates of 4 cover cropping treatments in a fully randomized block design, with a total of 24 plots containing 3 raised beds with a single line of squash each (see Supplemental Figure 1). The cover cropping treatments were Rye (*Secale cereale*) Cover Crop (RCC), Chemically Terminated Rye Cover Crop (CT-RCC), Plastic Cover (PC) and Bare Soil (BS). Specifically, for 2016 and 2017 growing season, the RCC treatment consisted of fall rye (cv Gauthier) seeded at a rate of 250kg/ha on September 14^th^ 2015 and September 16^th^ 2016 and rolled to the ground the following spring by crimping rye with a 3-sectional roller crimper(I&J Mfg., PA, USA) on June 13^th^ 2016/2017; the CT-RCC treatment was the same as RCC except rye was killed with an herbicide (glyphosate (Roundup, WeatherMaxMD, Bayer, Canada) at a rate of 2.16 kg a.e. ha^-1^) before the rye crimping; the PC treatment consisted in the application of an agricultural plastic mulch over each raised bed within plots; and the BS treatment was a bare soil mound lane that did not receive any cover cropping treatment. To monitor treatment effects on pathogen populations, we inoculated squash seeds with a rifampicin-resistant *P. syringae* strain prior to direct seeding into raised beds. The strain pathogenicity was confirmed by hypersensitive reaction (HR) testing in tobacco leaves and seed inoculation was validated by growing on-field control plants at the border of each treatment plot.

### Microbial collection, DNA extraction and sequencing

Microbial communities of the phyllosphere were collected from squash at three different times each growing season in 2016 (12 July,1 August and 1 September), and 2017 (12 June, 31 July and 5 September), as defined as *Early, Mid and Late* season. Each sample consisted of a mix of young and old leaves harvested from the squash canopy by clipping an average of 16.8±8.3 g and 20.7±4.8 g of leaves for years 2016 and 2017 respectively (see Supplemental Table 3) from an individual plant into sterile sample bags (SCR-7012-ID, Innovation Diagnostics Inc., Blainville, Canada) with surface-sterilized shears. Replicate samples (3 per plot) were collected for a total of 72 samples per sampling date (3 samples x 4 treatments x 6 replicates). Microbial cells were then gathered by washing each leaf sample using with 110 ml of saline buffer [10% NaCl] and using a homogeniser blender (Stomacher® 400, Seward, UK) for 30 sec at 250 rpm.

A volume of 1ml of wash solution from each sample was placed on King’s B (KB) medium with cycloheximide (50mg/l) (C7698, Sigma Aldrich, Oakville, CA) and rifampicin (50mg/l) (R3501, Sigma-Aldrich, Oakville, CA), allowing us to estimate *P. syringae* population size by counting colony forming units CFU after 4 days of growth at 28 C°. Specifically, pathogen counts were calculated as:

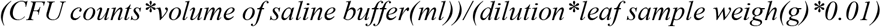

The remaining 100 ml wash solution was divided into two 50ml Falcon tubes; one was centrifuged at 11,500 × g for 20 min and the other at 4,500 × g for 20 min. The aqueous phase was removed from both tubes and the pellet in the Falcon tube centrifuged at 4,500 × g was frozen at -80°C. The DNA of the remaining pellet was extracted using MoBio PowerSoil DNA extraction kits (CA-11011-418, VWR, Mont-Royal, CA) and stored at -20°C for future processing. Amplicon libraries were prepared for Illumina sequencing using PCR targeting the V5–V6 region of the bacterial 16S rRNA gene using cyanobacteria-excluding primers [16S primers 799F-1115R] (29,30) to exclude chloroplast DNA (for a 16S amplicon structure overview, see supplemental Figure 2). The 25 microliter PCR reactions consisted of 5 μL 5x HF buffer (Thermo Scientific, Waltham, MA, USA), 0.75 μL DMSO, 0.5 μL dNTPs (10 mM each), 0.25 μL Phusion Hot Start II polymerase (Thermo Scientific), 1 μL each primer (5 μM), 1 μL of genomic DNA, and 15.5 μL molecular-grade water (IDT, Coralville, IA, USA). We included, onto the libraries, a negative control (1µl of sterile water; IDT, Coralville, IA, USA) as well as a positive control (1µl of *P. syringae* DNA) was included in each 96 well library plate. Libraries were checked on Agarose gels (2%), normalized with SequalPrep kit (A1051001, Life Technologies, Burlington, CA) on a Gilson robot (<u>Middleton</u>, WI, USA) and sequenced on a MiSeq (Illumina, San Diego, CA, USA). For each ear, samples were randomly assigned between two sequencing runs, representing a total of 4 runs.

### Sequence analysis

Sequencing adaptors were removed with the bbduk tool from bbmap (38.86, https://sourceforge.net/projects/bbmap/), with the following parameters: ktrim=r k=23 mink=11 hdist=1 tpe tbo (31). Sequences were thereafter demultiplexed allowing one mismatch on the barcode sequence with deMulMe (https://github.com/RemiMaglione/genomicScript/tree/master/deMulMe). Sequence barcodes were removed with cutadapt 2.10 in paired-end mode (32). In total we obtained 11,929,677 and 11,816,138 demultiplexed paired-end sequences, for 2016 and 2017 respectively. All subsequent data processing and computations were done with DADA2 1.12.1 (33) in R 3.6.0 (RC Team, 2013) and graphs were produced using the ggplot2 3.2.1 package (36) Sequences were trimmed and quality filtered with *filterAndTrim* with default parameters except: trimLeft = c(19, 26), truncLen = c(230, 210), maxEE=c(2,3); *trimleft* was set to remove both PCR primers and barcodes used for libraries preparation, *truncLen* and *maxEE* were set to yield filtered sequences with a quality around a phred score of 30. Amplicon sequence variants (ASVs) were constructed from filtered sequences with the following set of built-in DADA2 functions and their default parameters except as mentioned: *dada* in pseudo-pooling mode, *mergePairs* with minOverlap = 30, *collapseNoMismatch* with minOverlap = 240, *removeBimeraDenovo* with method=“pooled”. ASVs were then taxonomically annotated by *assignTaxonomy* with SILVA version 128 database (37,38). For 2016 and 2017 data, the DADA2 pipeline yielded a mean of 10,484,056 filtered paired-end sequences used to identify 7,604 ASVs, which represent an average of 29,652 sequences and 165 ASVs per sample.

A preliminary evaluation of positive and negative control samples was performed with a principal component analysis ordination of a distance matrix obtained with centred log ratio (clr) transformation of the original community matrix (39). Since control samples would be lost by excluding samples with very few sequences, the clr transformation allowed us to keep all the samples while identifying outlier samples. Since the control samples were distinct compositionally from the squash samples (see Supplemental Figure 3), they were removed for all further analyses.

### Data analysis

#### *P. syringae* abundance analysis on squash leaves

Since no significant differences in *P. syringae* abundances among experimental blocks were observed in 2016, therefore, *P. syringae* count differences between treatments were evaluated with a linear model of treatment effects on *P. syringae* abundance. In 2017, effects of experimental blocks on *P. syringae* abundances were significant and were thus integrated in a mixed linear model of treatment effects on abundances as a random effect. Effect of treatment was estimated with a TukeyHSD post-hoc test performed on the above-mentioned model.

#### Squash fruit health and marketability at harvest

We quantified squash fruit health and marketability by harvesting fruit within a 10m x 10m area within each plot. Fruit health and marketability was determined with 4 categories of *P. syringae* symptoms based on the visually estimated proportion of fruit affected by the considered symptoms: *P*.*syringae* symptoms outside of the fruits, *P*.*syringae* symptoms that penetrate the fruits, *P*.*syringae* symptoms that let a scare at the surface of the fruits and *P*.*syringae* symptoms that generate squash rot. Marketability has been assessed from each categories of *P. syringae* symptoms where more than 1% of fruit affected in at least one category prevent marketability. Healthy fruit was defined as a squash fruit with no *P. syringae* symptoms. Thus, marketability and fruit health were binomially distributed and their differences among treatments were evaluated with generalized mixed linear model, where blocking effect was integrated as random variable, for both years, except marketability in 2016 where blocking effect was not significant and treatment effect was modeled with generalized linear model. Effects of treatment on fruit health and marketability was estimated with a TukeyHSD post-hoc test (using the *glht* function of *multcomp* R package) performed on the above-mentioned model.

#### Effect of cover cropping treatments on bacterial community diversity

Diversity analyses were performed using the R package phyloseq 1.30.0 (40), picante 1.8.1 (41), and vegan 2.5-6 (34). To evaluate the effect of treatments on community diversity on squash leaves, samples were randomly rarefied to 5000 sequences per sample: this threshold was chosen to preserve the maximum number of samples with a sufficient quantity of ASVs to capture the majority of the diversity in each sample (see Supplemental Figures 4 and 5). For all diversity analyses, rarefactions and their subsequent analyses were repeated 1000 times but no qualitative differences were observed between iterations, and so we report here the results of a single random rarefaction of the data. The uniformity of relative abundance distributions of ASVs (alpha diversity) was assessed with the Shannon index (42). The effect of treatment on alpha diversity was evaluated with a post-hoc test (TukeyHSD) of a linear model (alpha diversity as a function of treatment). Variation in bacterial community structure among samples was quantified with the Bray-Curtis index (43). Major gradients in community composition were evaluated with nonmetric multidimensional scaling (NMDS) ordination of weighted Bray-Curtis distances among samples. We partitioned the variance in phyllosphere bacterial community structure explained by sampling date and treatment using generalized mixed model and permutational ANOVA analysis of the variance in Bray-Curtis distances. We tested sample clusters of two compositionally distinct groups of treatment with a least squares comparison between treatment and the two NMDS axis scores, with the emmeans v1.4.8 R package (44).

#### Differential abundance analysis of ASVs

Differential abundance analysis of ASVs among treatments were performed with DeSeq2 3.11 (45). The ASV matrix was filtered using the CoDaSeq R package 0.99.4 (46), with the *codaSeq*.*filter* function with the following parameters: min.reads=1000 (minimum reads per sample), min.prop=0.00001 (minimum proportional abundance of a read in any sample), min.occurrence=0.005 (minimum fraction of non-zero reads for each variable in all samples). Then, zero counts from the filtered matrix were substituted with the Bayesian-multiplicative replacement of zero counts approach included in the *cmultRepl* function from the zCompositions R package (47). Since DeSeq2 takes non-zero positive integers as input, we adjusted the values of the zero-replaced matrix (*mv*) so that the lowest value is equal to 1 with: *mv*+(1-min(*mv*)). DeSeq2 analysis was executed with parameters recommended for single-cell analysis that better fit data with a zero-inflated negative binomial distribution such as our modified community matrix. We tested for differential abundance by contrasting ASV abundances across all six possible treatment comparisons: Rye Cover Crop versus Chemically Terminated-Rye Cover Crop, Rye Cover Crop versus Plastic Cover, Rye Cover Crop versus Bare Soil, Chemically Terminated-Rye Cover Crop versus Plastic Cover, Chemically Terminated-Rye Cover Crop versus Bare Soil and Plastic Cover versus Bare Soil. We used the following model: *design = ∼ block + treatment* and the blocking random variable was controlled through the *reduced* parameter. Only contrasts with *adjusted P-value<=*0.01 and *log_2_-fold-change* ≥ 1 were considered to be significantly differentially abundant.

## Results

### Cover cropping reduced *P. syringae* abundance on squash leaves and improved fruit health and marketability

We found that *P. syringae* was less abundant on the leaves of squash grown with rye cover crops (Figure 1) and harvested squash fruits was more marketable and healthier with rye treatments (Table 1). In 2016, *P. syringae* CFU counts were significantly lower for the Rye Cover Crop treatment compared to Plastic Cover and Bare Soil treatments during the *Early* season (Tukey HSD post-hoc on linear model; Figure 1). There were no significant differences among treatments during *Mid* and *Late* season sampling in 2016. On the other hand, in 2017, *P. syringae* CFU counts were significantly lower during the *Early* season for both rye cover crop treatments (Rye Cover Crop and Chemically-Terminated Rye Cover Crop) compared to Plastic Cover and Bare Soil treatments (Tukey HSD post-hoc test on linear mixed model; Figure 1). No *P. syringae* colonies were retrieved at *Mid* season on the squash leaves grown with rye cover crops, and pathogen populations were lower for Chemically-Terminated Rye Cover Crop as compared to Plastic Cover and Bare Soil treatments. Finally, pathogen CFUs were significantly lower between Rye Cover Crop and Bare Soil at *Late* season sampling. Thus, *P. syringae* populations were significantly lower with rye cover cropping during the entire 2017 growing season. Taken together, P. syringae populations showed the greatest reduction in the rye cover crop treatments early in the growing season. This finding is consistently supported across the two years of the experiment (see Supplemental Table 1 and 2). Moreover, fruits health and marketability were significantly different across all year of harvest (p<0.05). Indeed, marketability with Rye Cover Crop was significantly different from Bare soil and Plastic Cover with an average marketability increase of, as compare to Bare Soil, 13% in 2016 and 6.17% in 2017, and as compare to Plastic Cover, 8% in 2016 and 4.33% in 2017 (Table 1). Fruit health enjoy an increase of 14% in 2016 with Rye Cover Crops treatment as compare to Bare Soil and 7.17% in 2017 with Chemically-Terminated Rye Cover Crop treatment as compare to Bare Soil (Table 1). No further significant differences were overserved for fruits health and marketability in both year of harvest.

**Figure 1:**
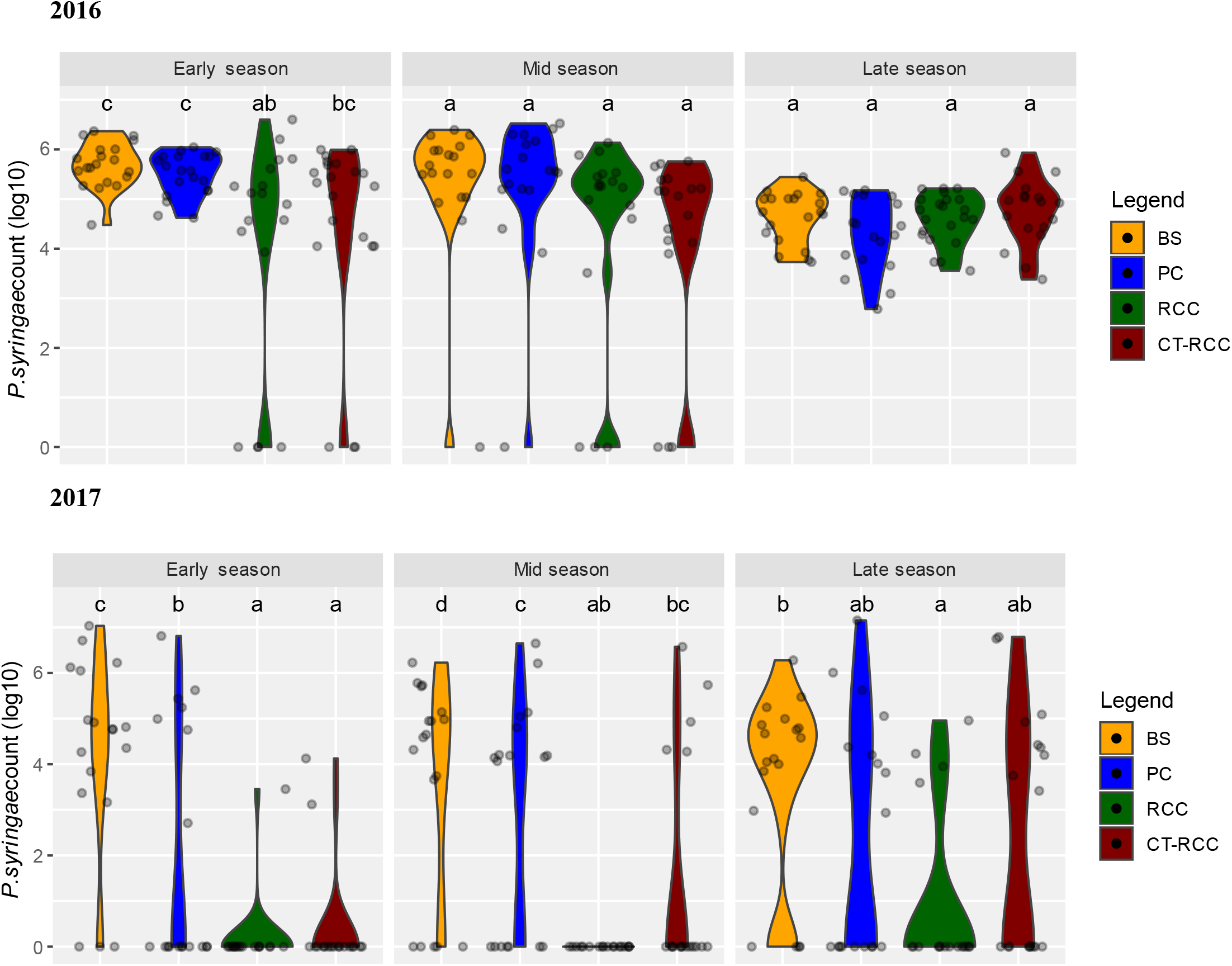
*P. syringae* population of squash leaves for different cover cropping practices during 2016 and 2017. Squash pathogen population sizes were estimated based on CFU count from bacterial culture of each leaves sample retrieved from 4 cropping treatments: Bare Soil (BS), Plastic Cover (PC), Rye Cover Crop (RCC) and Chemically Terminated Rye Cover Crop (CT-RCC). Different letters represent significantly different treatments (p<0.05) from a post-hoc test (TukeyHSD) of a linear model (*P. syringae* as a function of treatment) in 2016 or a linear mixed model (*P. syringae* as a function of treatment (fixed effect) and block (random effect) in 2017

**Table 1:** Proportion of squash fruit (mean +-standard deviation) with no *P. syringae* symptoms and marketable fruits with no damage for the two-growing seasons 2016 and 2017. Differences among treatments were tested using Tukey’s honestly significant difference (HSD) test, based on a generalized mixed model with General Linear Hypotheses provided by glht function of multcomp package. Treatments that do not share a letter were significantly different according to the Tukey HSD test (p<0.05). Since random effect of blocking variable for *Proportion of marketable squash fruit with no damages for 2016 growing season* was not significant, TukeyHSD has been evaluated this year on generalized linear model with TukeyHSD function of stats package

| Year | *Plastic Cover* | *Rye Cover Crop* | *Chemically-Terminated Rye Cover Crop* | *Bare Soil* |
| --- | --- | --- | --- | --- |
| **Proportion of squash fruit without *P. syringae* symptoms (%)** |  |  |  |  |
| 2016 | 56.8 ± 6.7 **bc** | 63.2 ± 7.7 **ab** | 59.7 ± 10.4 **bc** | 49.2 ± 9.3 **c** |
| 2017 | 79.3 ± 8.8 **bc** | 90.8 ± 6.9 **bc** | 87 ± 8.3 **ab** | 79.8 ± 8.4 **c** |
| **Proportion of marketable squash fruit with no damages (%)** |  |  |  |  |
| 2016 | 79.5 ± 5.4 **c** | 87.5 ± 5.8 **ab** | 81.3 ± 3.4 **bc** | 74.5 ± 7.3 **c** |
| 2017 | 91.17 ± 7.1 **c** | 95.5 ± 3.4 **ab** | 94.2 ± 3 **bc** | 89.3 ± 3 **c** |

### Phyllosphere microbial communities differed between sampling dates and treatments

Cover cropping treatments influenced bacterial community composition on squash leaves. Treatments also affect bacterial diversity and richness (see Supplemental Analysis 1 with Supplemental Figures 6 &7). A nonmetric multidimensional scaling ordination of the overall community distance matrix suggests that squash phyllosphere samples clustered by sampling dates in both years (see Supplemental Figure 8); sampling date accounted for 28% (R^2^ = 0.28, p<0.001) and 11% (R^2^ = 0.11, p<0.001) of community compositional variation between samples for 2016 and 2017 respectively (PERMANOVA on Bray-Curtis distances among samples). Because there was an interaction between sampling date and treatments (PERMANOVA on Bray-Curtis distances among samples; sampling date * cover crop treatment interaction P<0.001 both years), and sampling date accounted for the majority of the effect, we thus analyzed the effect of treatments on communities separately for each date in order to summarize these complex effects.

### Effect of treatments on community diversity

Bacterial community alpha diversity was significantly different among treatments for several sampling dates (linear model; Shannon diversity vs. cover cropping treatment, P<0.05; Figure 2). Bacterial community alpha diversity was higher for both rye treatments as compared to bare soil and plastic treatments in 2016 in the *Early* season sampling (Tukey HSD on linear model; Figure 2). No further differences were observed between treatments at the other sampling dates in 2016. Although both rye cover cropping practices resulted in significantly lower Shannon diversity in *Mid* season sampling as compared to Bare Soil, their alpha diversity was higher as compared to plastic treatments in *Early* season sampling. No further differences were observed between treatments in late season sampling of 2017. Taken together alpha diversity increased early in the growing season for both rye cover cropping treatments as compared to bare soil and plastic treatments of 2016 or to the plastic treatment of 2017.

**Figure 2:**
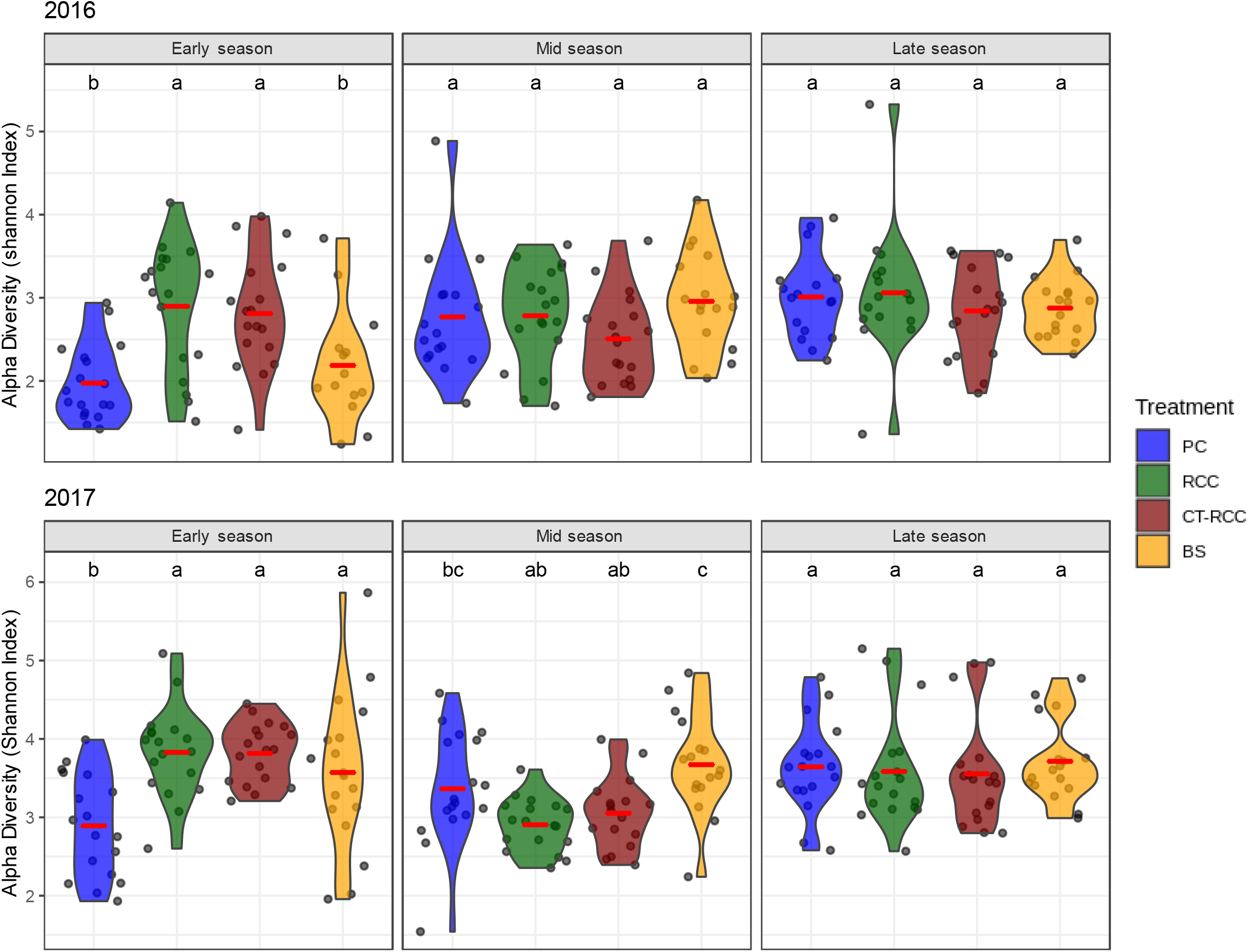
Violin plot of alpha Diversity (Shannon index) for each treatment and each sampling date during the growing season of years 2016 and 2017. Horizontal red line represents the mean distribution. Blue: PC (Plastic Cover), green: RCC (Rye Cover Crop), red: CT-RCC (Chemically Terminated Rye Cover Crop) and yellow: BS (Bare Soil). Different letter represents significantly different treatments (p<0.05) from a post-hoc test (TukeyHSD) of a linear model (alpha diversity as a function of treatment)

### Cover cropping treatments influence squash phyllosphere communities

Community composition varied among cover cropping treatments for each sampling date of 2016 and 2017 (PERMANOVA on Bray-Curtis distances for each sampling date; cover cropping effect P<0.001). Moreover, distances between treatment clusters for the ordination (Figure 3) suggests that treatment effects were more important for *Early* season sampling as compared to the other sampling dates. Differences in community composition among treatments were more pronounced in *Early* (PERMANOVA on Bray-Curtis distances; effect of cover cropping treatment P_2016 & 2017_<0.001; R^2^_2016_ = 0.24, R^2^_2017_ =0.31) rather than *Mid* (P_2016 & 2017_<0.001; R^2^_2016_ = 0.14, R^2^_2017_ =0.30) and *Late* season sampling (P_2016 & 2017_< 0.001; R^2^_2016_ = 0.07, R^2^_2017_ = 0.16) (Figure 3).

**Figure 3:**
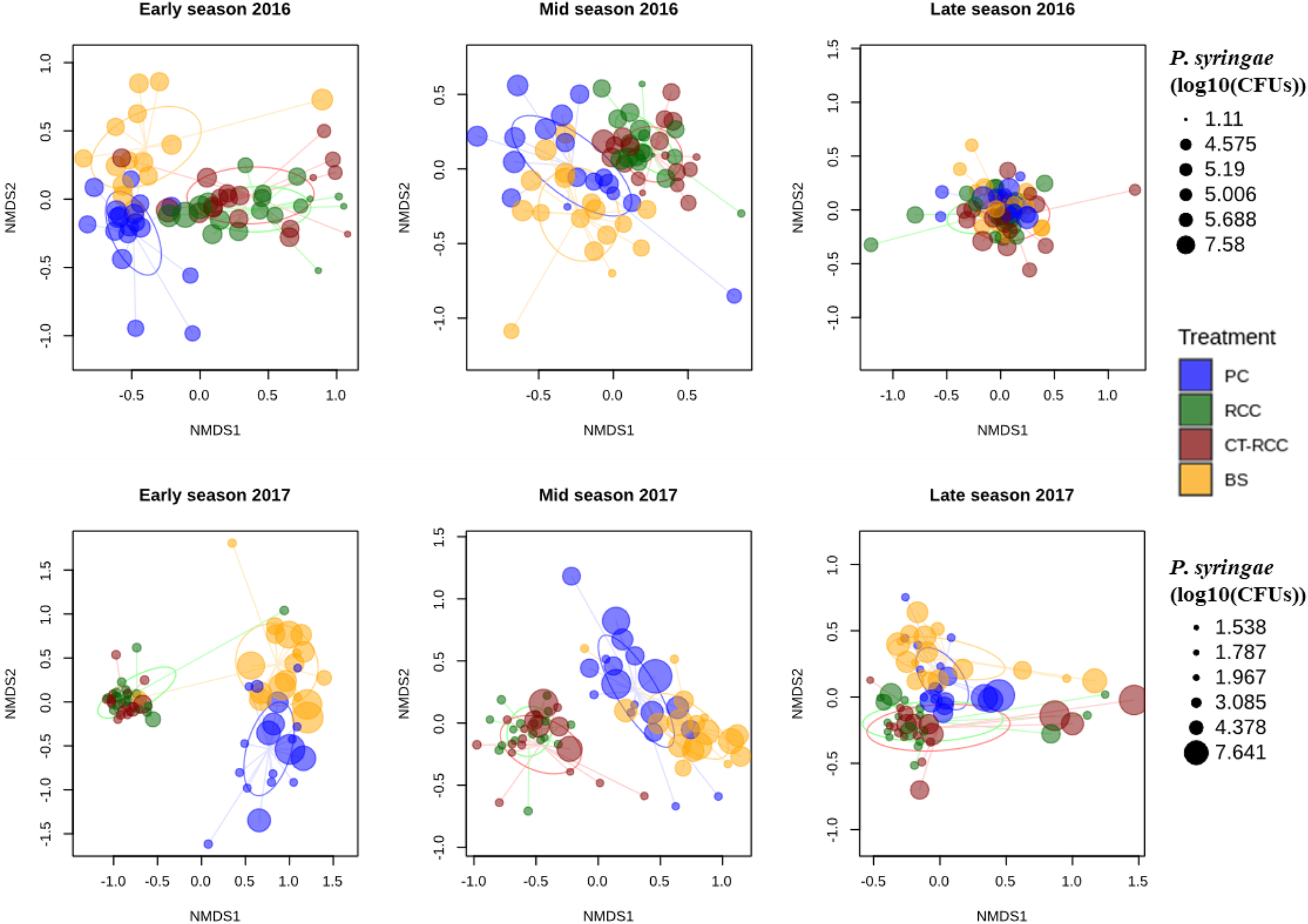
Non-metric multidimensional scaling (NMDS) ordination of bacterial community composition in squash phyllosphere samples from different cover cropping treatments in 2016 and 2017. Each point represents a phyllosphere community; symbol size indicates the abundance of *P. syringae* colony forming units (log10(CFUs)) in that sample; colors indicate the cover cropping treatment: blue: PC (Plastic Cover), green: RCC (Rye Cover Crop), red: CT-RCC (Chemically Terminated Rye Cover Crop) and yellow: BS (Bare Soil).

Ordination of samples based on community composition also indicated that samples clustered into two compositionally distinct groups: Rye and Chemically-Terminated Rye Cover Cropping, versus Plastic and Bare Soil (Figure 3). At each sampling date, sample scores on the first axis of ordination differed significantly between cover cropping treatments. Rye and Chemically-Terminated Rye were different from the Plastic and Bare Soil, but not different from each other (linear mixed model with blocking variable as random effect and Tukey HSD on linear mixed model; ordination axis scores versus cover cropping treatment; Supplemental Figure 9). Moreover, according to the Tukey HSD, these two compositionally distinct groups (Rye and Chemically-Terminated Rye versus Plastic and Bare Soil) are more different during the *Early* season as compared to every other sampling date. This difference remains, although it progressively decreases, throughout the growing season. However, plastic and bare soil seem to not share this behavior, and those treatments were different in *Mid* season in 2017 and were often separated along the second axis of the NMDS ordination (Figure 3; estimated marginal means test of ordination axis scores; Supplemental Figure 9). Taken together, both rye treatments have a similar effect on the phyllosphere bacterial community structure as compared to plastic and bare soil. This effect was present during the entire growth period but was more pronounced early in the growing season.

### Differentially abundant taxa among treatments at each sampling date: *Sphingomonas* and *Methylobacterium* were more abundant with cover crop treatments

An analysis of differential abundance of ASVs among treatments and sampling dates identified several ASVs that were more abundant in certain treatments and at certain times. As mentioned previously, the cover crop effect on taxa abundance was analyzed separately for each sampling date because the abundance of *P. syringae* and the squash phyllosphere communities were also influenced by the date when the sampling occurred, differences being more important earlier in the season. To identify ASVs that were strongly associated with different cover cropping systems we took the top differentially abundant ASVs with the highest log_2_-fold change in abundance for each treatment comparison (2017: Figure 4, 2016: Supplemental Figure 10). Different cover cropping treatments had several differ rentially abundant ASVs with log_2_-fold changes in abundance between treatments ranging from -10.2 to 9.7. Overall, the contrasts of Rye versus Chemically-Terminated Rye, and Plastic versus Bare Soil consistently had fewer and weaker differentially abundant ASVs in comparison with the contrasts between these two groups. In 2016, ASVs that were significantly more abundant for Rye and Chemically-Terminated Rye Cover Crop treatments included those annotated at the genus level as *Rhizobium, Pseudomonas* and *Saccharibacillus* during the *Early* season, and *Chryseobacterium* and *Sphingomonas* during the *Mid* season. Conversely, ASVs that were significantly more abundant in Bare Soil and Plastic Cover treatments included those annotated as *Pseudoatrobacter* during the *Early* season, *Exiguobacterium* and *Pseudoatrobacter* during the *Mid* season, and *Deinococcus* during the *Late* season. In 2017, ASVs that were significantly more abundant for Rye and Chemically-Terminated Rye Cover Crop treatments included those annotated as the genera *Sphingomonas, Methylobacterium or Hymenobactrium* during the *Early* season, *Sphingomonas, Methylobacterium, Aureimonas* and *Microbacterium* during the *Mid* season, and *Chryseobacterium* and *Rhizobacterium* during the *Late* season. On the other hand, ASVs that were significantly more abundant in Bare Soil and Plastic Cover treatments included, *Massila* and *Exiguobacterium* at *Early* season, *Massila, Exiguobacterium, Hymenobacter, Deinococcus* and *Pseudoatrobacter* at *Mid* season and *Deinococcus* and *Microbacterium* at *Late* season.

**Figure 4:**
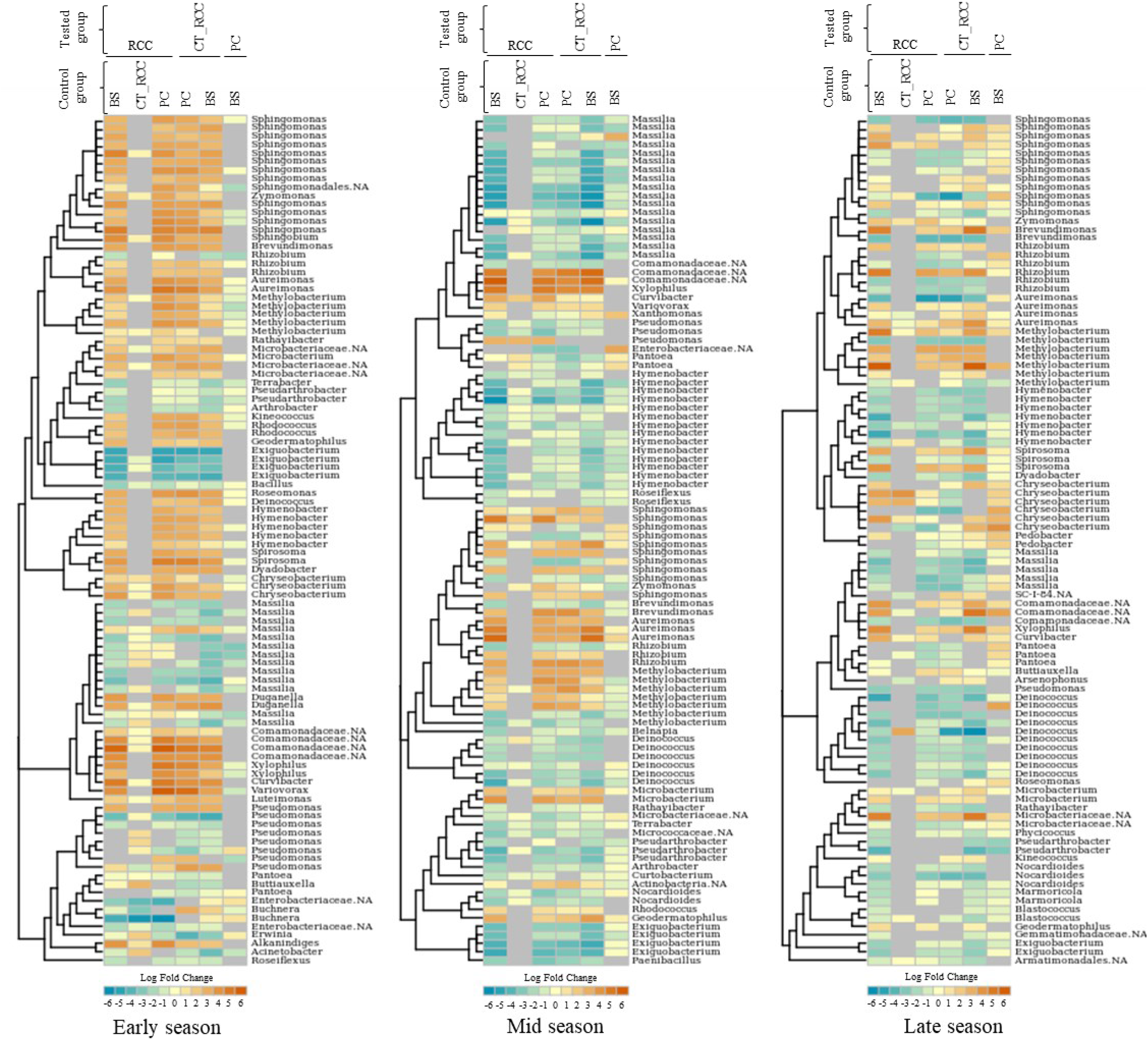
log2-fold change (LFC) heatmap of most differentially abundant ASV from DeSeq2 analysis for each sampling date of the 2017 samples. For each panel, left track is the phylogenetic tree from pynast alignment of ASVs sequence while right track is the corresponding taxonomic name at the Genus rank. Each heatmap column is a different contrast between two treatment mentioned in header as followed: above name is the “tested” treatment whereas the below one is the “control” treatment meaning that a positive LFC value represent an ASV more abundant for the tested treatment. Grey color represents no LFC for the ASV. Each number on the bottom LFC colour scale represents a level of LFC. Tested treatment: PC (Plastic Cover), RCC (Rye Cover Crop), CT-RCC (Chemically Terminated Rye Cover Crop) and yellow: BS (Bare Soil).

## Discussion

Rye cover cropping reduced the abundance of *P. syringae* on squash leaves, improved the health and marketability of fruit, and shaped phyllosphere bacterial community composition and diversity. The greatest effect of cover cropping on both the phyllosphere community and *P. syringae* abundance was observed early in the growing season. *P. syringae* begins life on leaves as an epiphyte but then must colonize host tissue through stomata or wounds (48). Disease severity could be lowered if the early establishment and survival of *P. syringae* is jeopardized. Taken together our results suggest that cover cropping treatments induce both shifts in phyllosphere microbiota and a protective effect against *P. syringae*, by reducing leaf pathogen abundance and fruit symptoms, with a critical early-season window of strongest microbiota changes associated with a protective effect against epiphytic pathogen colonization. We inoculated seeds with *P. syringae* at the time of planting, but in real situations, the effect of cover crops on pathogen populations will be a function of temporal variation in seed and soil pathogen and microbiota reservoirs.

We found that both Rye and Chemically-Terminated Rye Cover Cropping treatments induced a strong shift in the squash phyllosphere microbiota, leading to a distinct community composition in comparison with Bare Soil and Plastic Cover treatments. The largest difference in leaf bacterial community composition associated with rye cover crop was observed early in the growing season. Copeland et al. (2015) reported homogenisation of community structure over time; early in the growing season, leaf microbiota was more diverse and soil was a strong driver of phyllosphere microbiome. Moreover, shifts in microbial community composition are likely driven by changes in environmental conditions as well as shifts in sources of bacterial migration to the phyllosphere. Cover cropping can directly modify soil abiotic properties such as temperature and moisture and chemical properties (50). Cover cropping also likely influences bacterial dispersal sources both by promoting colonization by bacteria living on the cover crops themselves, as well as through their effects on dispersal from different potential sources such as soils (51), water splash (52) and insects (53,54). Taken together, we hypothesize that such local environmental shifts modify bacterial migration to the phyllosphere early in the growing season.

In addition to the effects of cover cropping on phyllosphere microbial communities, cover cropping treatments also likely influenced environmental conditions, which may explain part of their protective effects against *P. syringae*. Cover crops influence humidity and temperature (10). We observed that soil moisture and temperature varied among cover cropping treatments; soil moisture was higher under rye cover crops and plastic cover relative to bare soils, and temperatures were elevated under plastic cover relative to other treatments (Maglione et al., personal communication). Rye is also known to have allelopathic properties (55), which are also known to influence soil microbiota (56). Moreover, rye degradation can lead to decreases in soil pH (57) and improve weed control (58). All of these effects of cover cropping on the abiotic and biotic environment could influence early pathogen development, and interact with shifts in phyllosphere microbiota to amplify the potential protective effects of cover crops.

Previous studies have reported a protective effect against pathogens by phyllosphere microbial diversity *per se* (59), for example where increasing *Sphingomonas* diversity on leaves increased protection against *P. syringae* (21). In our study, there was not strong evidence for an effect of alpha diversity on its own to explain the protective effect of cover cropping against *P. syringae*; there were no overall differences in alpha diversity of phyllosphere bacteria among cover cropping treatments, although rye cover cropping did increase diversity in the early season. To properly test for a protective effect of phyllosphere diversity against *P. syringae*, future studies that directly manipulate diversity while keeping other factors constant will be required, but our results suggest that it was the composition of bacterial communities and not the diversity of the community *per se* that could explain the protective effects of cover cropping treatments.

Our results support the hypothesis that rye cover crops could protect against *P. syringae by* promoting the establishment of potential competitors or plant growth promoting bacteria (PGPB) on the leaf surface. Rye cover cropping led to increases in the abundance of numerous bacterial phyllosphere taxa (60). This included several ASVs belonging to the genus *Sphingomonas*, which were more abundant with rye cover cropping, especially early in the growing season. Previous studies have demonstrated that *Sphingomonas* strains protect *Arabidopsis* against *P. syringae* in a controlled environment (21). Our findings provide field-based evidence suggestive that the *Sphingomonas* clade is a potential PGPB. We also found many other taxa preferentially associated with squash under rye cover cropping treatments, including ASVs belonging to the genus *Methylobacterium* that is a phyllosphere-associated clade (61) known to be an important PGPB in agriculture (62), and ASVs belonging to the genus *Pseudomonas*, which has been shown to be an antagonist of the pathogens *Erwinia* (63), Tobacco Necrosis Virus (64), and *Botrytis cinereal* (65). Thus, rye cover crops appear to favor the establishment of plant beneficial bacteria in the phyllosphere. The potentially beneficial bacterial ASVs associated with rye cover crops that we have identified are candidates for exploration of microbiome engineering approaches to directly inoculate protective bacterial strains to protect crops against pathogens (66).

### Conclusion

In conclusion, here we have shown that rye cover cropping is a sustainable agriculture practise that protects squash against the pathogen *P. syringae*. We have also provided evidence of a strong effect of rye cover crops on bacterial communities of the squash phyllosphere. Leaf microbial communities shifts as well as protective effects against *P. syringae* epiphytic development were more striking early in the growing season, during the crucial period of early plant growth and leaf colonization by *P. syringae*. We identified numerous bacterial ASVs belonging to the genera *Sphingomonas, Methylobacterium* and *Pseudomonas* that were promoted by rye cover crops. Thus, cover crops offer a means of sustainable management of bacterial pathogens and a reservoir of potential biocontrol agents. Open questions that remain include understanding the sources of the bacterial populations that were promoted by cover cropping; did they colonize the phyllosphere directly from the rye cover crop, or did cover cropping indirectly select for beneficial bacteria by altering environmental conditions? To our knowledge, this is the first study to investigate how cover cropping practices impact phyllosphere bacterial communities, and our results show that rye cover cropping is a very promising sustainable agriculture practise that protect squash against pathogens and is potentially mediated by shifts in phyllosphere microbial communities.

## Availability of data and materials

The demultiplexed sequence data have been deposed as sequences read archive under the BioProject: PRJNA705113. The scripts used to perform analyses for the current study are available in a GitHub repository: https://github.com/RemiMaglione/Science-Communication/tree/main/Article/cover-crop-squash-phyllosphere-microbiota-2021

## Acknowledgements

We thank Katherine Bisaillon, Pierre-Olivier Hebert and Jean-Baptiste Leducq for their help in the field, in the lab, with analysis and for discussions.

