## Supplementary material for "Winter rye cover cropping changes squash (*Cucurbita pepo*) phyllosphere microbiota and reduces *Pseudomonas syringae* symptoms": Winter rye cover cropping changes squash (Cucurbita pepo) phyllosphere microbiota and reduces Pseudomonas syringae symptoms Supplemental.pdf

### SUPPLEMENTAL MATERIALS

**Supplemental Table 1. Means of *P. syringae* colony forming units (CFUs) recovered from squash leaves grown in different cover cropping practices.** Means are reported as log10-tranformed number of colonies (± standard deviation) for the four cropping practices at the three-season sampling

| Mean CFUs (log10) ± standard deviation | | | | |
|---|---|---|---|---|
| **Year/** *Season* | ***Plastic Cover*** | ***Rye Cover Crop*** | ***Chemically-Terminated Rye Cover Crop*** | ***Bare Soil*** |
| **2016** | | | | |
| *Early* | 4.06 ± 2.33 | 4.46 ± 1.97 | 5.49 ± 0.43 | 5.69 ± 0.45 |
| *Mid* | 4.38 ± 2.1 | 4.17 ± 1.99 | 5.29 ± 1.49 | 5.33 ± 1.42 |
| *Late* | 4.6 ± 0.5 | 4.75 ± 0.67 | 4.29 ± 0.74 | 4.64 ± 0.52 |
| **2017** | | | | |
| *Early* | 0.19 ± 0.81 | 0.4 ± 1.18 | 1.98 ± 2.65 | 4.19 ± 2.2 |
| *Mid* | 0 ± 0 | 1.44 ± 2.43 | 2.93 ± 2.5 | 3.58 ± 2.38 |
| *Late* | 0.93 ± 1.81 | 2.43 ± 2.63 | 2.4 ± 2.62 | 3.59 ± 2.1 |

**Supplemental Table 2: *P. syringae* CFUs (recovered from squash leaves) ratio between different cover cropping practices at each sampling date during 2016 and 2017.** Ratios of significantly different treatments according to Tukey HSD; *: p<0.05; **p<0.01; ***p<0.001 *italic value*: no *P. syringae* CFU were recovered at date 2 in 2017 for RCC treatments.

| **2016** | **CT-RCC/RCC** | **PC/RCC** | **BS/RCC** | **PC/CT-RCC** | **BS/CT-RCC** | **BS/PC** |
|---|---|---|---|---|---|---|
| *Early* | 2.51 | **26.9*** | **42.08*** | 10.74 | 16.8 | 1.56 |
| *Mid* | 0.61 | 8.05 | 8.99 | 13.19 | 14.74 | 1.12 |
| *Late* | 1.4 | 0.48 | 1.08 | 0.35 | 0.78 | 2.25 |
| **2017** | **CT-RCC/RCC** | **PC/RCC** | **BS/RCC** | **PC/CT-RCC** | **BS/CT-RCC** | **BS/PC** |
| *Early* | 1.62 | **60.71*** | **9871.12***** | **37.43*** | **6086.54***** | **162.59**** |
| *Mid* | *27.27* | *858.06**** | *3796.51**** | 31.47 | **139.22**** | 4.42 |
| *Late* | 31.56 | 29.48 | **459.35**** | 0.93 | 14.56 | 15.58 |

**Supplemental Table 3: Mean weight (g) of squash leaves samples for each sampling date during years 2016 and 2017.** Mean weights (g) are reported with their standard deviation

| Season | 2016 | 2017 |
|---|---|---|
| *Early* | 16.91±5.34 | 20.38±4.50 |
| *Mid* | 11.49±10.86 | 19.21±3.74 |
| *Late* | 21.62±4.14 | 22.44±5.54 |
| All | 16.81±8.32 | 20.67±4.82 |

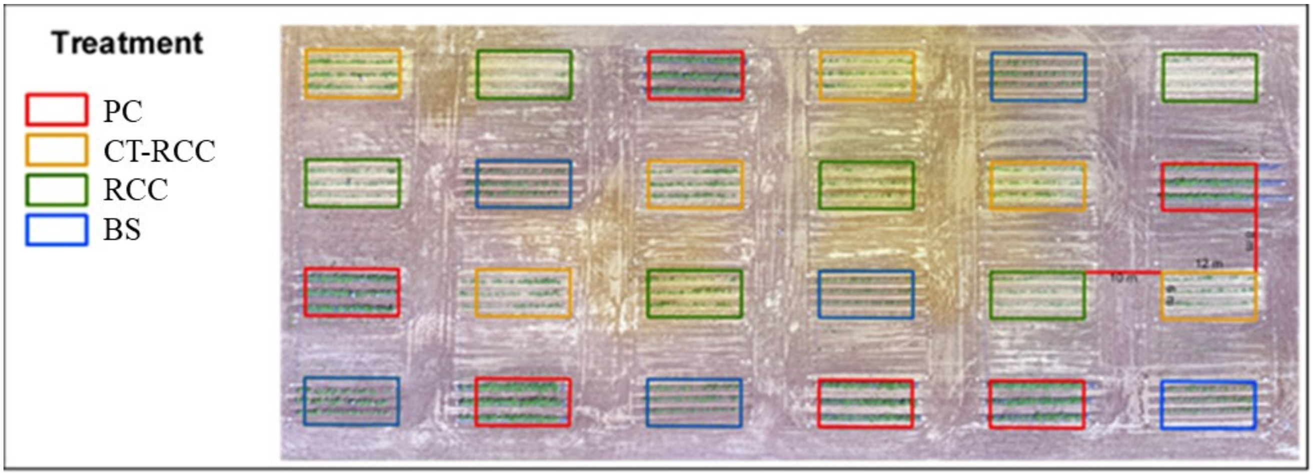


**Supplemental Figure 1**: **Aerial photo of the experimental field with cover cropping treatments indicated for each experimental plot.** Each experimental plot is 12m long by 6m wide. Plots within the same column are separated by 8-10m between plots of the same row. Each column defines a block of four randomized treatments. Each treatment was applied on 3 raised-bed lines of squash within each experimental plot. Treatment : PC (Plastic Cover), RCC (Rye Cover Crop), CT-RCC (Chemically Terminated Rye Cover Crop) and yellow: BS (Bare Soil).

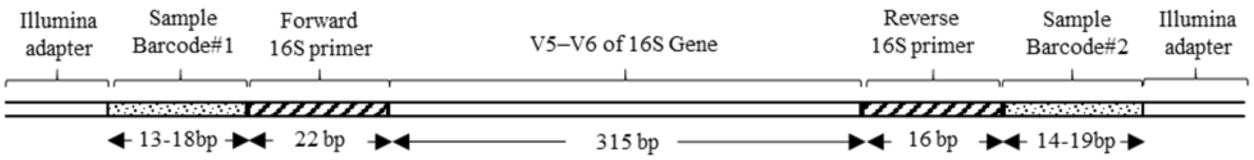


**Supplemental Figure 2**: **Structural overview of 16S amplicons.** Each amplicon owns a dual indexing (combination of Barcode#1 and #2) for each sample. Barcode length are variable between samples.

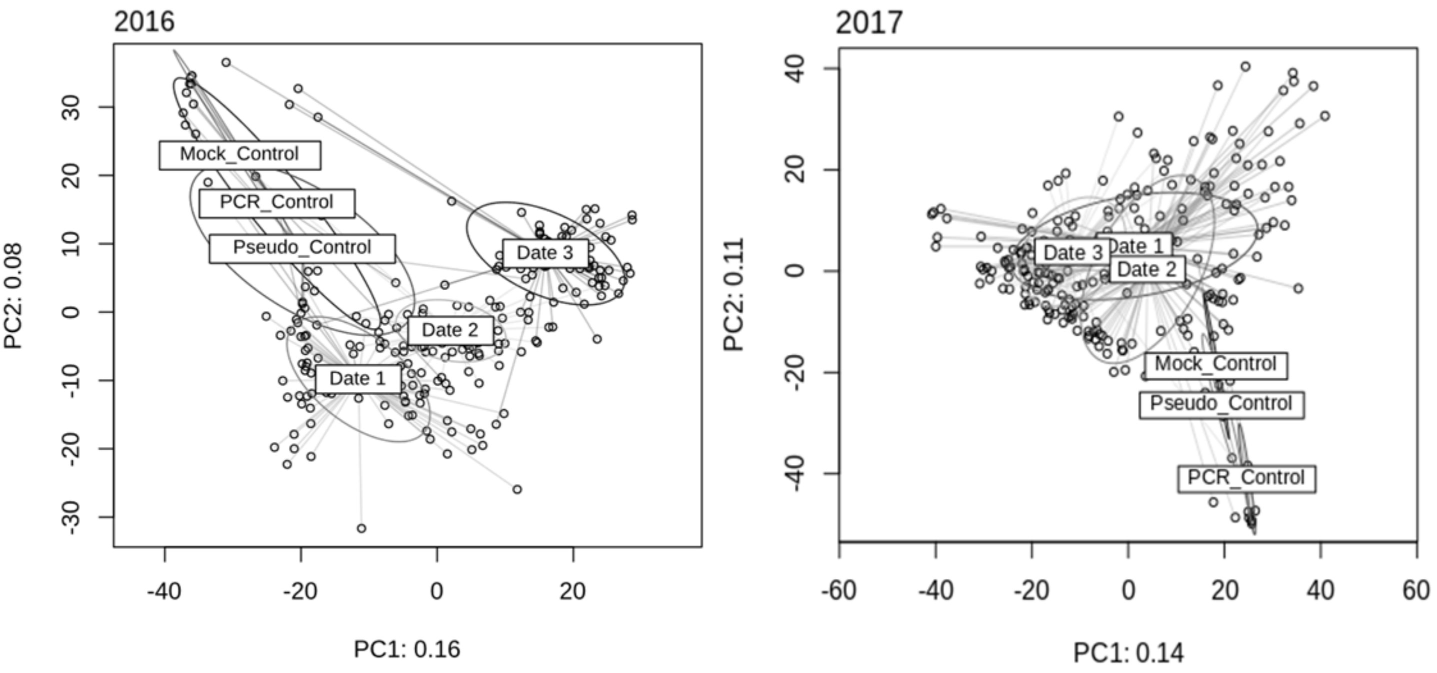


**Supplemental Figure 3: Principal component analysis ordination among phyllosphere samples and control samples for the growing season 2016 and 2017.** Ellipses represent standard deviation around samples from each category. Control samples, including positive controls Mock_Control (known mix of 5 bacterial isolate) and Pseudo_Control (*P. syringae* isolate) and negative control PCR_Control (water), are distinct compositionally from the phyllosphere samples. Corresponding sampling dates are as follows : *Early*=Date 1, *Mid*=Date 2, *Late*=Date 3.

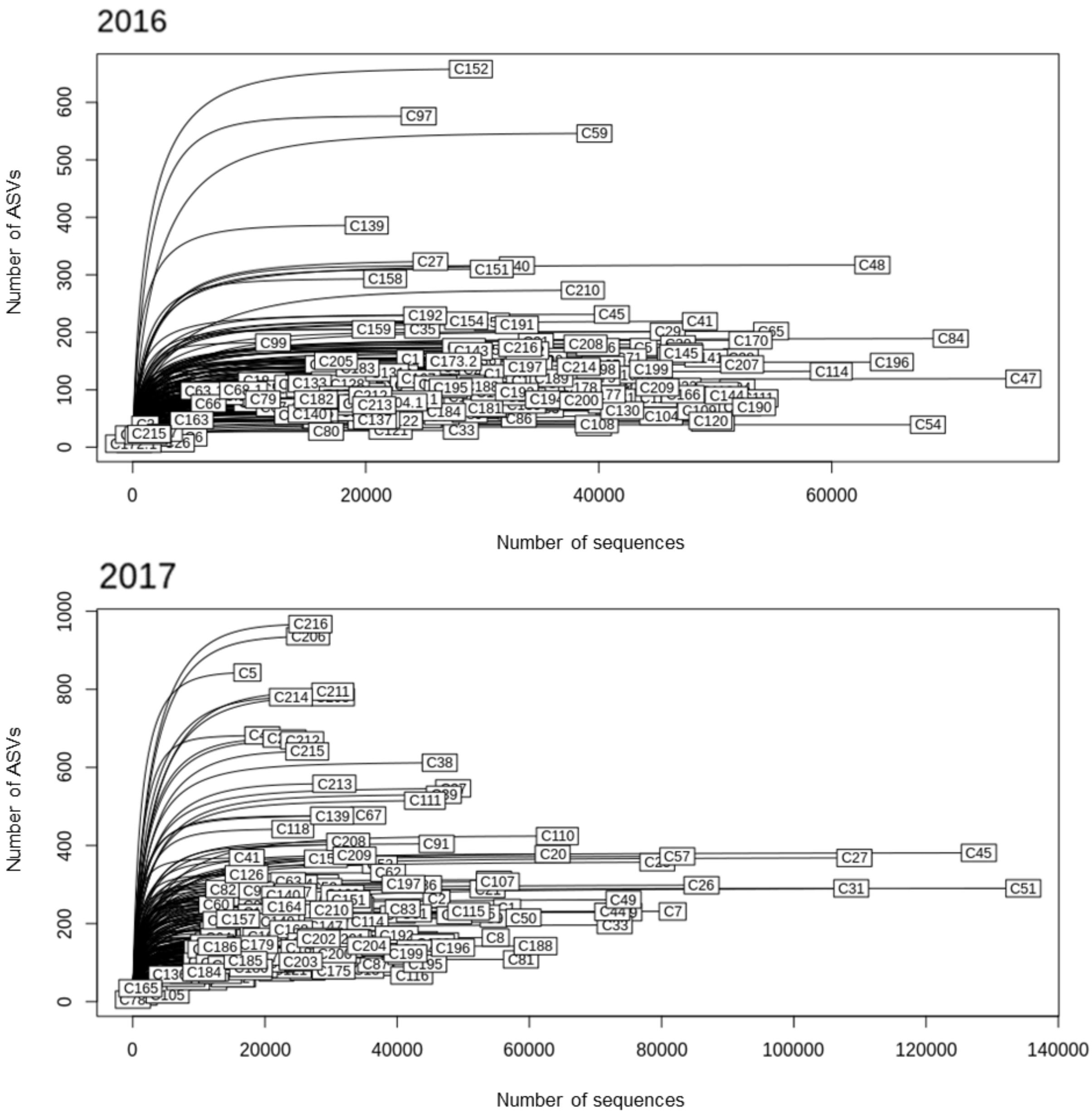


**Supplemental Figure 4. Rarefaction curves (ASVs versus number of sequences/sample) for each squash sample of growing season 2016 and 2017.** Each label represents a different sample. The flat section of each curve indicates that enough sequences has been reach to capture the majority of ASVs for a given sample.

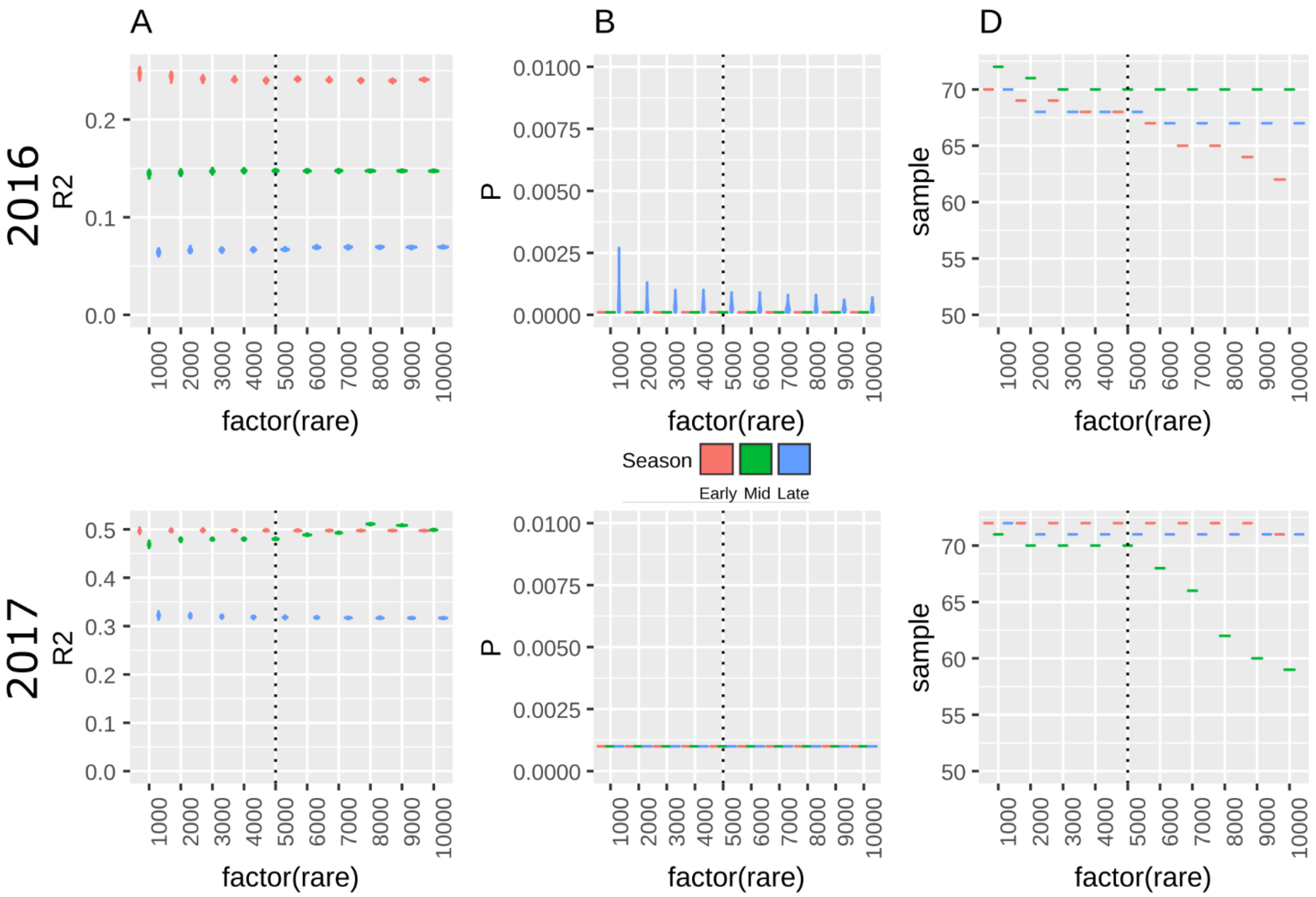


**Supplemental Figure 5: effect of rarefaction level over analysis and results.** 100 iteration of rarefaction levels, from 1000 to 10000 random sampling per sample, where tested to evaluate their effects on: $R^2$ (A) and p-value (B) of the permANOVA (per date effect of treatment on community structure), and remaining sample after rarefaction (C). The vertical dotted line represents the chosen rarefaction level for our study.

**Supplemental Analysis 1: Effect of treatments on overall community diversity**

Cover cropping treatments influenced the diversity of the bacterial community on squash leaves as measured by the Shannon diversity of rarefied bacterial communities (supplemental Figure 6). During the growing season, the community diversity was marginally significantly higher for the rye treatment in 2016 (TukeyHSD post-hoc test performed on linear model of community diversity * treatment; $p=0.0565$) and significantly higher for the bare soil (TukeyHSD post-hoc test performed on linear mixed model of community diversity ~ treatment with block as the random effect $p= 0.0332$) in 2017 as compare to the other treatments whereas overall treatment effect was marginally significant (liner model of community diversity ~ treatment; 2016: $p=0.086$, 2017: $p= 0.053$). Moreover, community diversity was significantly different between sampling dates (linear model of community diversity ~ sampling date; 2016, $p> 0.001$, 2017: $p= 0.003$). While this diversity was lower for sampling date 2 as compare to sampling date 1 (TukeyHSD post-hoc test performed on the above-mentioned linear model; $p= 0.031$) and date 3 ($p= 0.003$) in year 2017 (no significant differences between date 1 and 3), it was higher for both sampling dates 2 (TukeyHSD post-hoc test performed on the above-mentioned linear mixed model; $p= 0.045$) and 3 ($p< 0.001$) than the sampling date 1 in year 2016 (no significant difference between date 2 and 3).

The total richness of ASVs in bacterial communities also differed among treatments and dates (supplemental Figure 7). No significant ASVs number differences between treatments were observed during the 2016 overall growing season (liner model of community richness ~ treatment; $p=0.201$), however it was statistically different in 2017 ($p< 0.001$) with more ASVs in the bare soil ($p< 0.001$) as compared to other treatments. Unlike the alpha diversity of 2016, the ASVs numbers were different between sampling dates (liner model of community richness ~ sampling date; $p= 0.009$) and significantly lower at date 2 (TukeyHSD post-hoc test performed on the above-mentioned linear model; $p= 0.009$). On the contrary, as the overall alpha diversity, the ASVs number was still different in the growing season in 2017, with less ASVs at sampling date 2 as compared to dates 1 ($p< 0.001$) and 3 ($p= 0.067$).

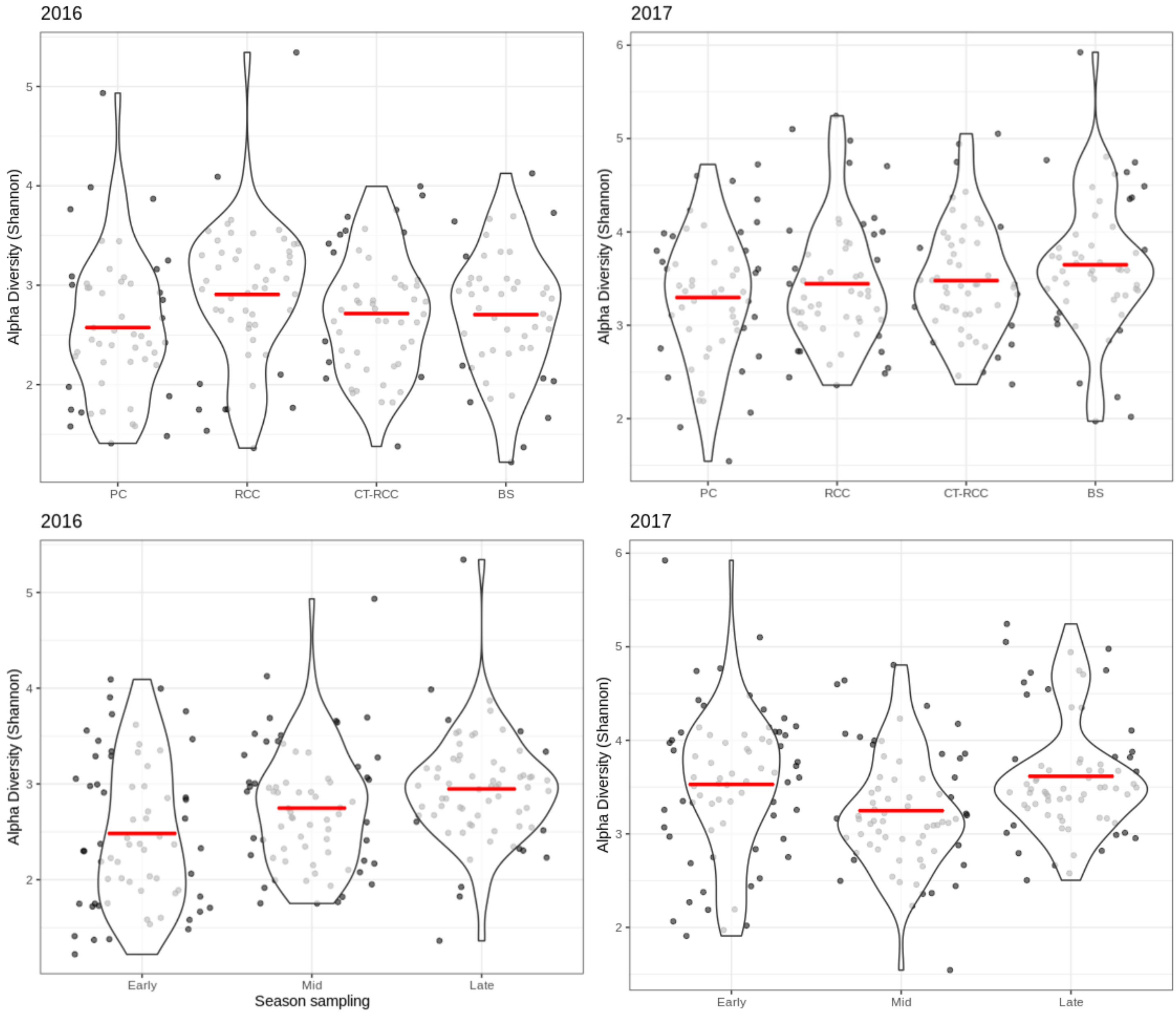


**Supplemental Figure 6: Violin plot of alpha diversity (Shannon diversity) for each treatment and sampling dates (*Early*, *Mid* or *Late* season) during the growing season of years 2016 and 2017.** Horizontal red lines represent the mean value. PC (Plastic Cover), RCC (Rye Cover Crop), CT-RCC (Chemically Terminated Rye Cover Crop) and yellow: BS (Bare Soil).

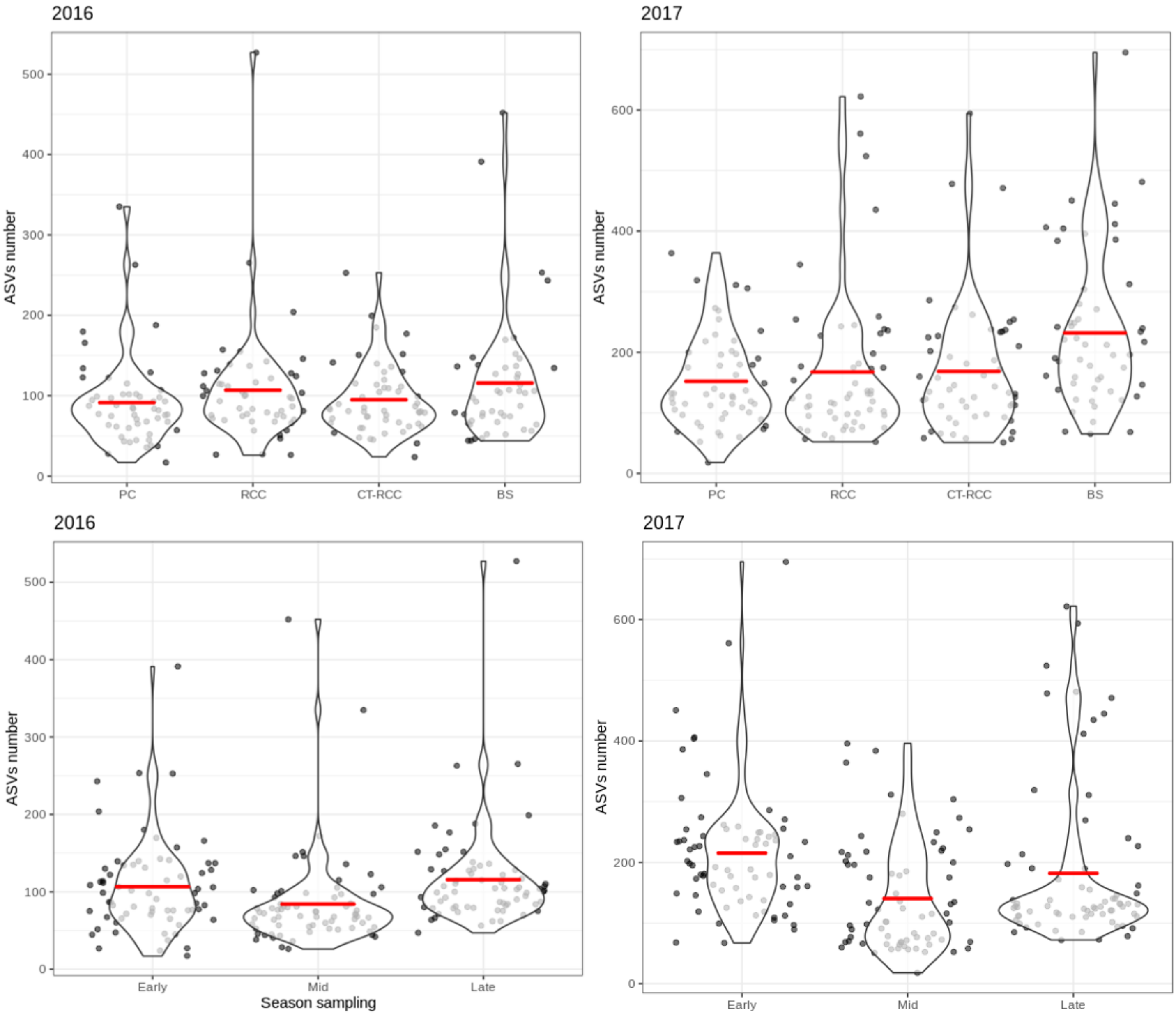


**Supplemental Figure 7: Violin plot of ASV richness at each treatment and sampling date during the growing season of years 2016 and 2017**. Horizontal red line represents the distribution mean. PC (Plastic Cover), RCC (Rye Cover Crop), CT-RCC (Chemically Terminated Rye Cover Crop) and yellow: BS (Bare Soil).

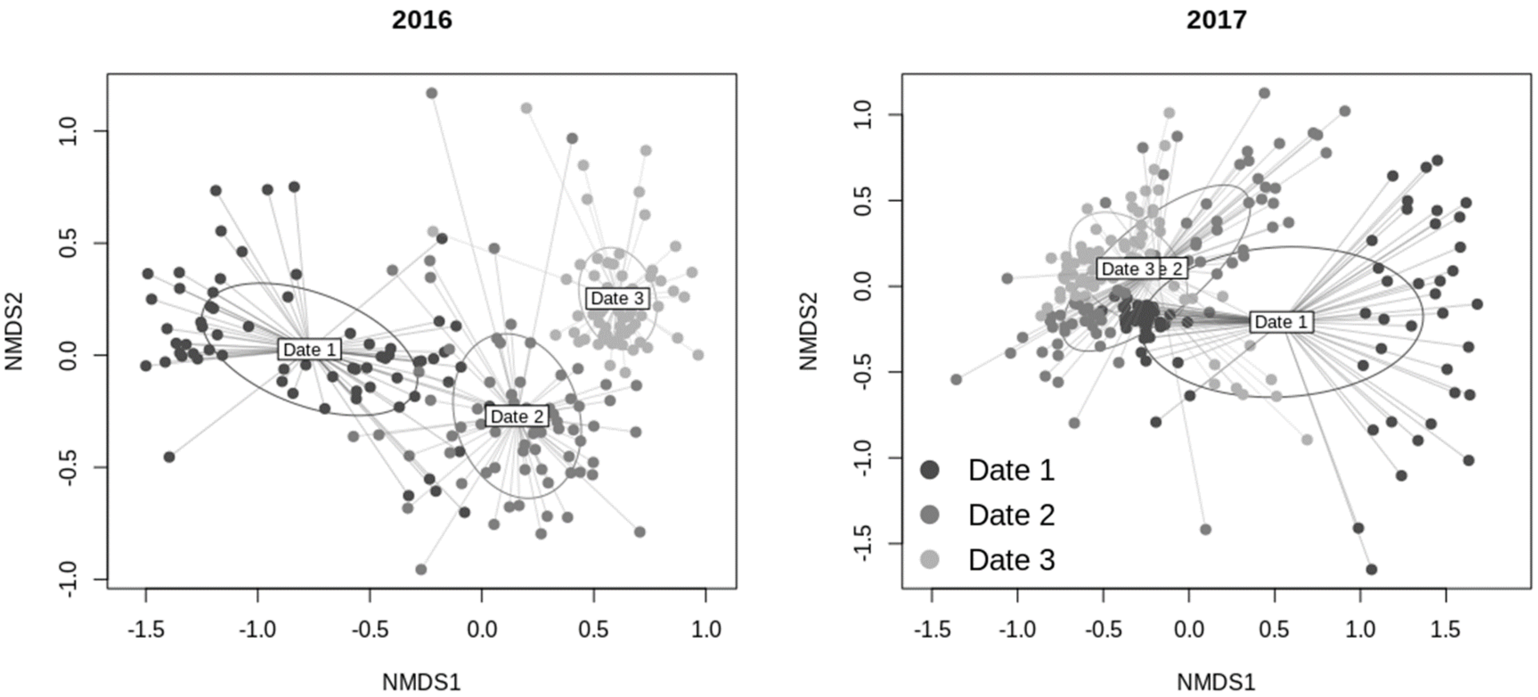


**Supplemental Figure 8**: ***ordination* of squash phyllosphere community during the growing season of 2016 and 2017 at the 3 sampling dates.** Corresponding season sampling dates are as follows: *Early*=Date 1, *Mid*=Date 2, *Late*=Date 3

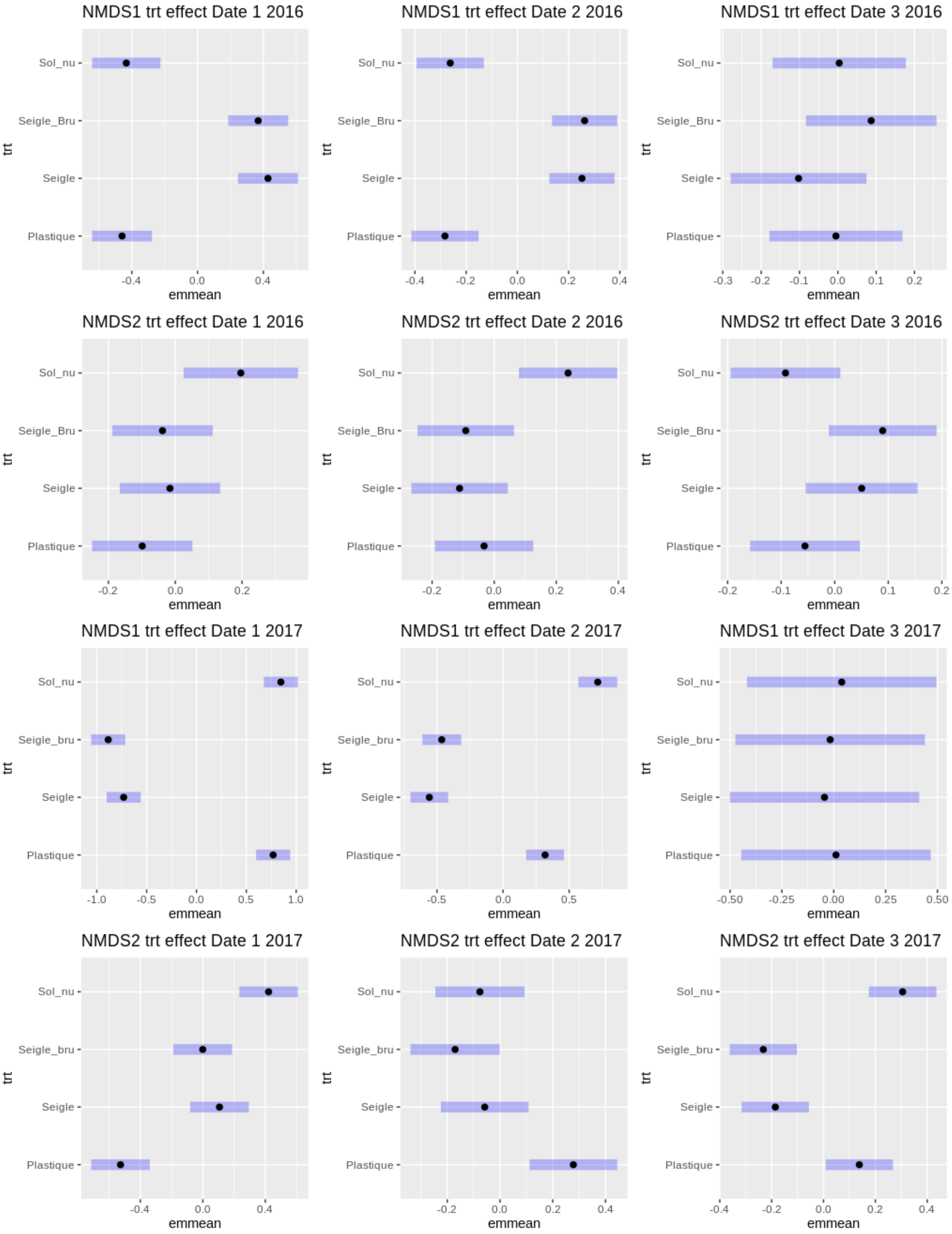


**Supplemental Figure 9**: least square plot of mixed model comparison between treatment and the 2 NMDS axis scores, at each sampling date of the growing season 2016 and 2017. Two overlapping horizontal bars indicating that the 2 treatments are not significantly different. The more distant the 2 bars, the greater the differences between the 2 associated treatments. Corresponding season sampling dates are as follows: *Early*=Date 1, *Mid*=Date 2, *Late*=Date 3

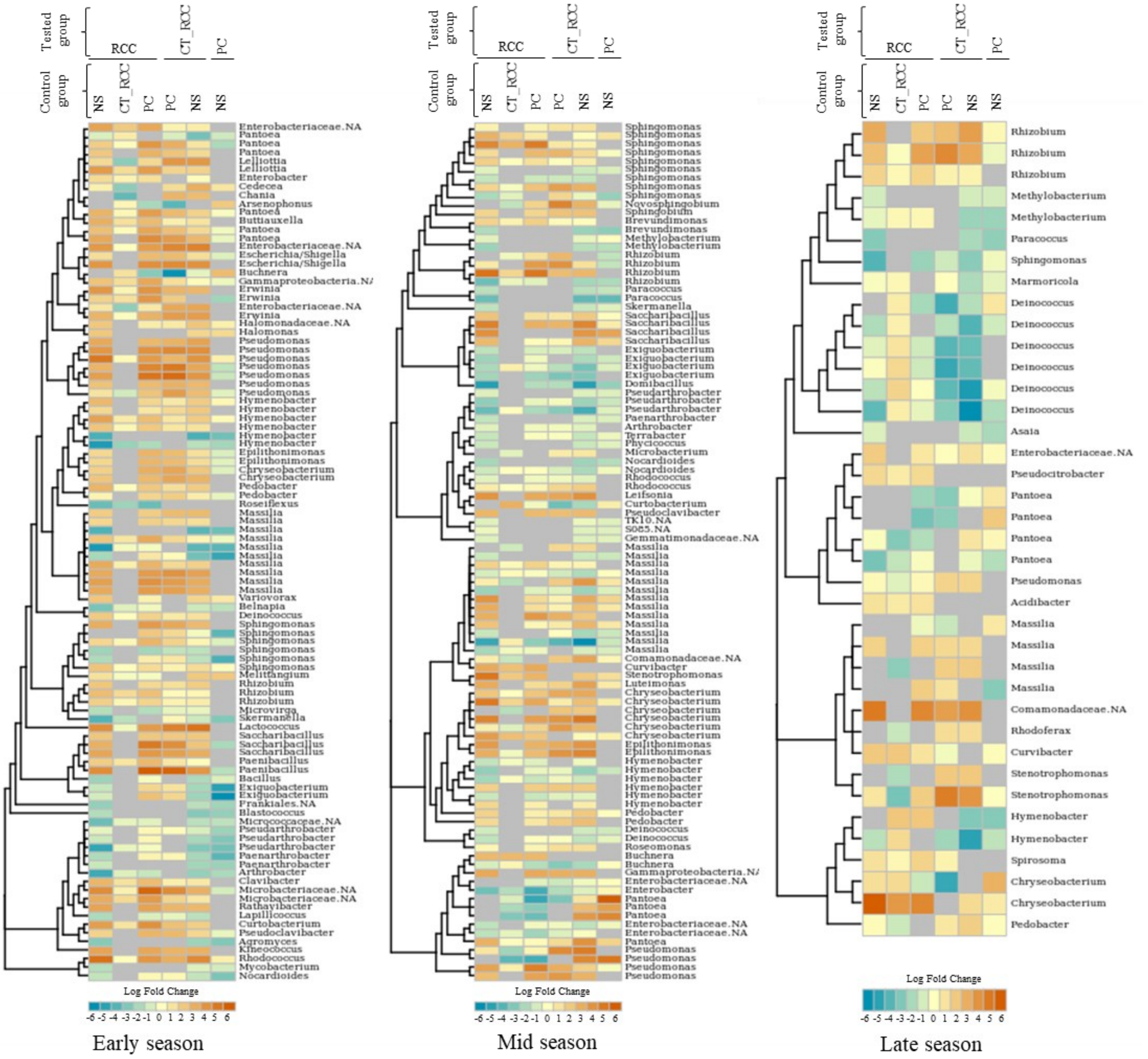


**Supplemental Figure 10**: **log2-fold change (LFC) heatmap of most differentially abundant ASV from DeSeq2 analysis for each sampling date of the 2016 samples**. For each panel, left track is the phylogenetic tree from pynast alignment of ASVs sequence while right track is the corresponding taxonomic name at the Genus rank. Each heatmap column is a different contrast between two treatments mentioned in the header as follows: above name is the "tested" treatment whereas the below one is the "control" treatment meaning that positive an LFC value represent an ASV more abundant for the tested treatment. Grey color represents no LFC for the ASV. Tested treatment: PC (Plastic Cover), RCC (Rye Cover Crop), CT-RCC (Chemically Terminated Rye Cover Crop) and yellow: BS (Bare Soil).
